# Clustering and temporal organization of sleep spindles underlie motor memory consolidation

**DOI:** 10.1101/2022.11.29.518200

**Authors:** Arnaud Boutin, Ella Gabitov, Basile Pinsard, Arnaud Boré, Julie Carrier, Julien Doyon

## Abstract

Sleep benefits motor memory consolidation, which is mediated by sleep spindle activity and associated memory reactivations during non-rapid eye movement (NREM) sleep. However, the particular role of NREM2 and NREM3 sleep spindles and the mechanisms triggering this memory consolidation process remain controversial. Here, simultaneous electroencephalographic and functional magnetic resonance imaging (EEG-fMRI) recordings were collected during night-time sleep following the learning of a motor sequence task. Adopting a time-based clustering approach, we provide evidence that spindles iteratively occur within clustered and temporally organized patterns during both NREM2 and NREM3 sleep. However, the clustering of spindles in trains is related to motor memory consolidation during NREM2 sleep only. Altogether, our findings suggest that spindles’ clustering and rhythmic occurrence during NREM2 sleep may serve as an intrinsic rhythmic sleep mechanism for the timed reactivation and subsequent consolidation of motor memories, through synchronized oscillatory activity within a subcortical-cortical network involved during learning.

## 1. Introduction

The repeated practice of a motor skill is crucial for its initial acquisition^1^. Yet, ample evidence from both animal and human studies indicate that following this early learning phase, the resulting motor memory trace is thought to be dynamically maintained during wakefulness and actively reprocessed (reactivated) during a subsequent sleep period^2–4^. These offline periods are believed to offer a privileged time window for the memory to be consolidated, a process whereby a newly acquired and relatively labile memory is transformed into an enhanced and more stable memory trace^5,6^. Indeed, a night of sleep (or even a daytime nap) has long been shown to play a crucial role in the strengthening and transformation of motor memory representations during consolidation, particularly for skills requiring the learning of a sequence of movements that is explicitly known to participants^7–12^. At the behavioral level, these changes can result in either the stabilization or spontaneous improvement in performance on the motor task during the post-sleep testing session (see ^4,13–15^ for systematic and meta-analysis reviews).

This memory-promoting effect of sleep in the consolidation of motor skills has been widely shown to rely on the repeated reactivation of newly encoded memories during non-rapid eye movement (NREM) sleep in both animal^16–18^ and humans^8,11^, and especially during NREM-stage2 (NREM2) sleep^7,19^. Growing evidence suggests that brief 0.3-2s bursts of sigma-like thalamocortical oscillations during NREM sleep, referred to as sleep spindle activity (~11-16 Hz), support such an offline covert reactivation of the memories to be consolidated^7,20–23^. Yet, the precise neurophysiological mechanism underlying such a spindle-related reactivation process in humans is still conjectural. On the one hand, spindles are known to undergo refractory periods of 3-6 s during which another spindle is unlikely to occur, hence limiting memory reactivations in time^13,23–26^. Such spindle refractoriness has been proposed as one of the possible mechanisms responsible for segregating memory reactivations and for protecting the reprocessing of memory traces without interference from unrelated information for a few seconds. On the other hand, our group has further suggested another possible mechanism based upon the rhythmic and timed occurrence of sleep spindles in clusters, also called “trains” (i.e., more than two successive spindles lasting less than 6 seconds in between), which would be relevant for the orderly repeated reprocessing of memory traces^13^.

Indeed, spectral dynamics of spindle rhythms during NREM sleep have been found to be cortically driven by infra-slow oscillatory rhythms^27–29^. More specifically, recent evidence shows that sleep spindles tend to cluster on a time scale of about 50 s (~0.02 Hz rhythm) during NREM sleep, with relative spindle-free periods in between during which some spindles may sporadically occur in isolation^20,27–29^. This NREM sleep infra-slow periodicity may thus constitute a critical neurophysiological regulatory mechanism providing transient temporal windows of optimal neural conditions for the reactivation and interference-free reprocessing of memory traces through spindle activity. Interestingly, the co-modulation of thalamocortical spindle and hippocampal sharp wave-ripples (SWR), which are linked to neural replay, has also been described over this infra-slow time scale^29^. Thus, beyond merely reflecting periodic spindle clustering, this ~0.02 Hz infra-slow sleep rhythm is further thought to offer cyclic periods of favorable conditions for the consolidation of hippocampus-dependent declarative memory^27^. However, the role of infra-slow fluctuations of sleep spindle activity in the consolidation of motor memories remains to be determined.

In the context of motor sequence learning (MSL), recent EEG-fMRI sleep studies have revealed that MSL consolidation relies on the synchronized local reprocessing of new information, time-locked to spindles, across segregated but inter-connected brain regions that were involved in the initial learning process *per se*, such as the hippocampus, striatum, thalamus and motor-related cortical areas^7,8,11,13^. Given the anatomical selective distribution and co-occurrence of spindles in the aforementioned brain regions^8,30,31^, recent work has reported increased functional connectivity within this network by showing enhanced hippocampal-cortical and striatal-cortical communication during sleep spindles^7,31–33^. More specifically, using deep brain EEG coherence analyses, Boutin and colleagues^7^ revealed that oscillatory synchrony in the spindle frequency band may reflect the cross-structural reactivation and communication between learning-related brain regions during sleep consolidation. However, it is still unknown whether such network-specific synchronized oscillatory activity during consolidation is related to sleep spindle clustering and rhythmicity. Hence, in the present study, we recorded simultaneous EEG-fMRI sleep data and conducted EEG deep brain coherence analyses to determine the functional contribution of sleep spindles, occurring either in train or in isolation, in the consolidation of a newly learned sequence of movements. Adopting a time-based clustering approach, we hypothesized that rhythmic occurrence of spindles in trains would confer favorable conditions for efficient reprocessing and consolidation of the memory trace within a functional network integrating the hippocampus, the striatum (the putamen in particular), the thalamus, and motor-related cortical circuitry^10,13,34^. In addition, considering the ubiquitousness of spindles over the cortex, we sought to determine whether the topographical expression (i.e., spatial distribution) of spindle clustering patterns may relate to the reprocessing of interrelated memory units during sleep. Hence, we expected sleep spindles to cluster preferentially locally over the cortical regions involved in motor sequence learning and consolidation, such as the pre-motor, sensorimotor, and parietal cortices^4,35,36^.

## 2. Results

### 2.1. Methods: overall experimental approach

Twenty participants were required to practice a motor sequence learning (MSL) task during training (14 blocks) and two retention sessions (1 block each) while lying supine in the MRI scanner. The MSL task required participants to perform an explicitly known 5-element finger movement sequence as rapidly and accurately as possible using a response pad. Each practice block was composed of twelve repetitions of the motor sequence (i.e., 60 key presses per block). Participants underwent the training session about 30 minutes before going to sleep. They were then administered two delayed retention tests, performed respectively 15 minutes (test) and 24 hours (retest) after the end of training. Immediately following the 15-min retention test, a simultaneous EEG-fMRI recording scan lasting a maximum of 2.25 hours took place while participants slept in the scanner. Following the EEG-fMRI sleep recordings, EEG electrodes were removed and participants were allowed to sleep for the rest of the night in the nearby sleep laboratory.

### 2.2. Behavioral performance

We quantified MSL performance by computing a score sensitive to both speed and accuracy. The performance score (PS) was calculated by dividing the number of accurately typed sequences by the time between key presses^7,37^. Mean PS during training and testing sessions are illustrated in Figure 1. Motor skill acquisition and consolidation were analyzed using a Block (Block 1, Block 14, Test, Retest) ANOVA with repeated measures on the factor Block. The analysis revealed a significant Block effect, *F*(3, 19) = 41.2, *p* < .01, *n*^2^_p_ = .68. Duncan’s multiple range test revealed significant within-session performance improvements from Block 1 to Block 14 (M_Block1_ = 200 and M_Block14_ = 290, *p* < .001), indicating that participants improved task performance during training. However, the post-hoc analysis failed to detect significant changes in performance from the test to the retest (M_Test_ = 302 and M_Retest_ = 284, *p* = .18), thus revealing a sleep-based stabilization rather than an enhancement of the consolidated memory (see ^38,39^ for reviews). At the group level, the absence of delayed performance gains over the retention interval may be due to the non-averaging of performance blocks during training and testing sessions, since block averaging is known to be a confounding factor that accounts for, and further exacerbates, the offline gains^14^.

**Figure 1.**
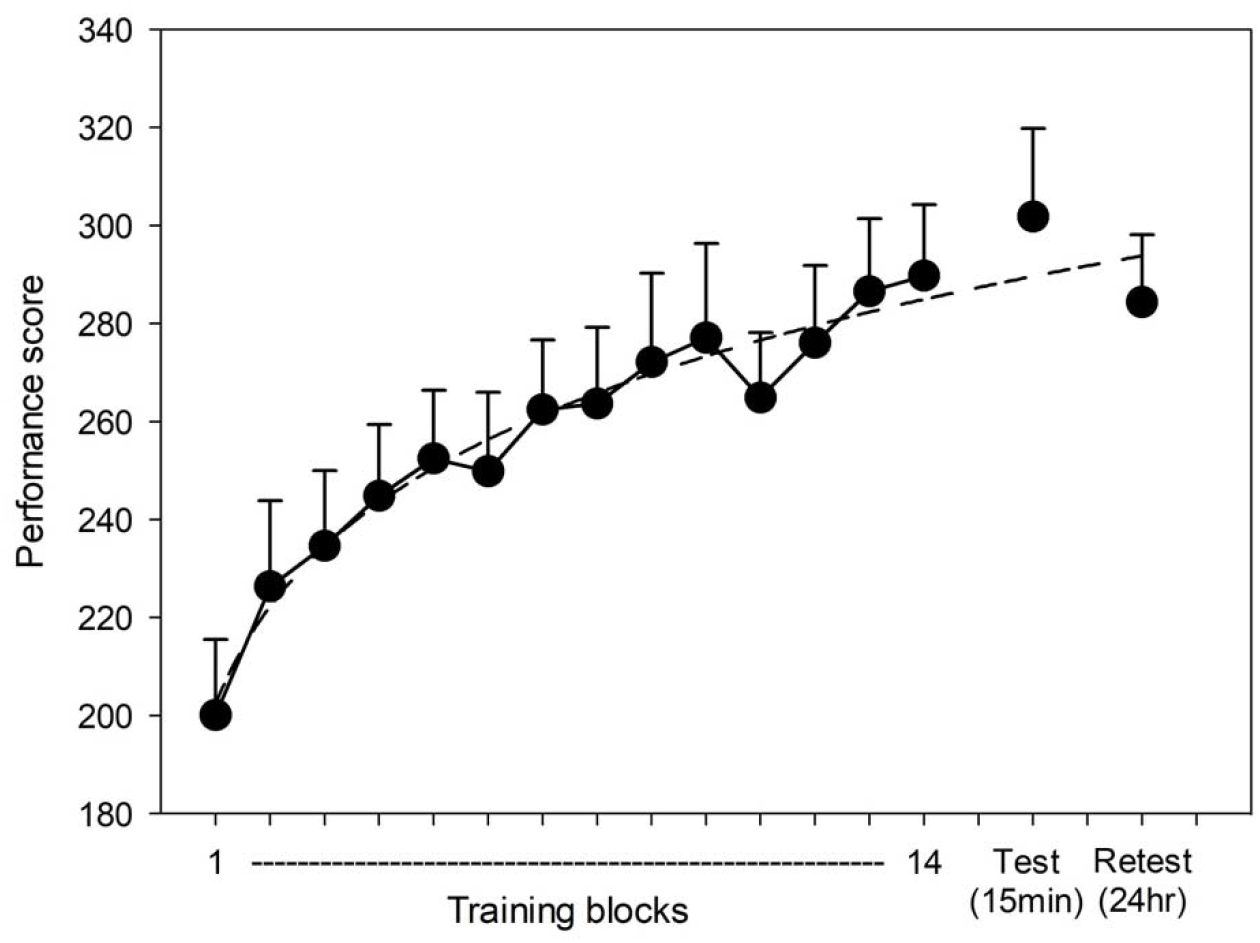
Behavioral results. Mean performance scores during training, delayed 15min test and 24hr retest blocks. A power-law function was fitted to the training data (dashed line). Error bars reflect the standard errors of the means.

### 2.3. Sleep spindle clustering and rhythmicity

#### Time-Frequency analysis

Figure 2A depicts the grand average time-frequency (TF) maps for spindle events occurring at scalp electrode Pz during NREM2 and NREM3 sleep periods, using epoch windows ranging from −75 s to +75 s around spindle onsets. TF analysis confirmed that spindles tend to cluster on a low-frequency time scale of about 41 ± 15 seconds during NREM2 sleep, and of about 54 ± 15 seconds during NREM3 sleep, when the analysis was restricted to NREM sleep periods with ITI lasting no longer than 180 seconds to rule out potential sub-infraslow periodic fluctuations of spindle-rich periods (which corresponds to the inclusion of 79.2 % of the full dataset). The ITI difference between NREM2 and NREM3 sleep reached significance, *t*(28) = −2.31, *p* = .023, *d* = −.86. Nonetheless, the group-averaged TF map revealed that spindle clustering predominates during NREM2 sleep, as illustrated by the long-range fluctuations of power increases in the 11-16 Hz spindle frequency range, in comparison to more erratic spectral dynamics of spindle rhythms during NREM3 sleep.

**Figure 2.**
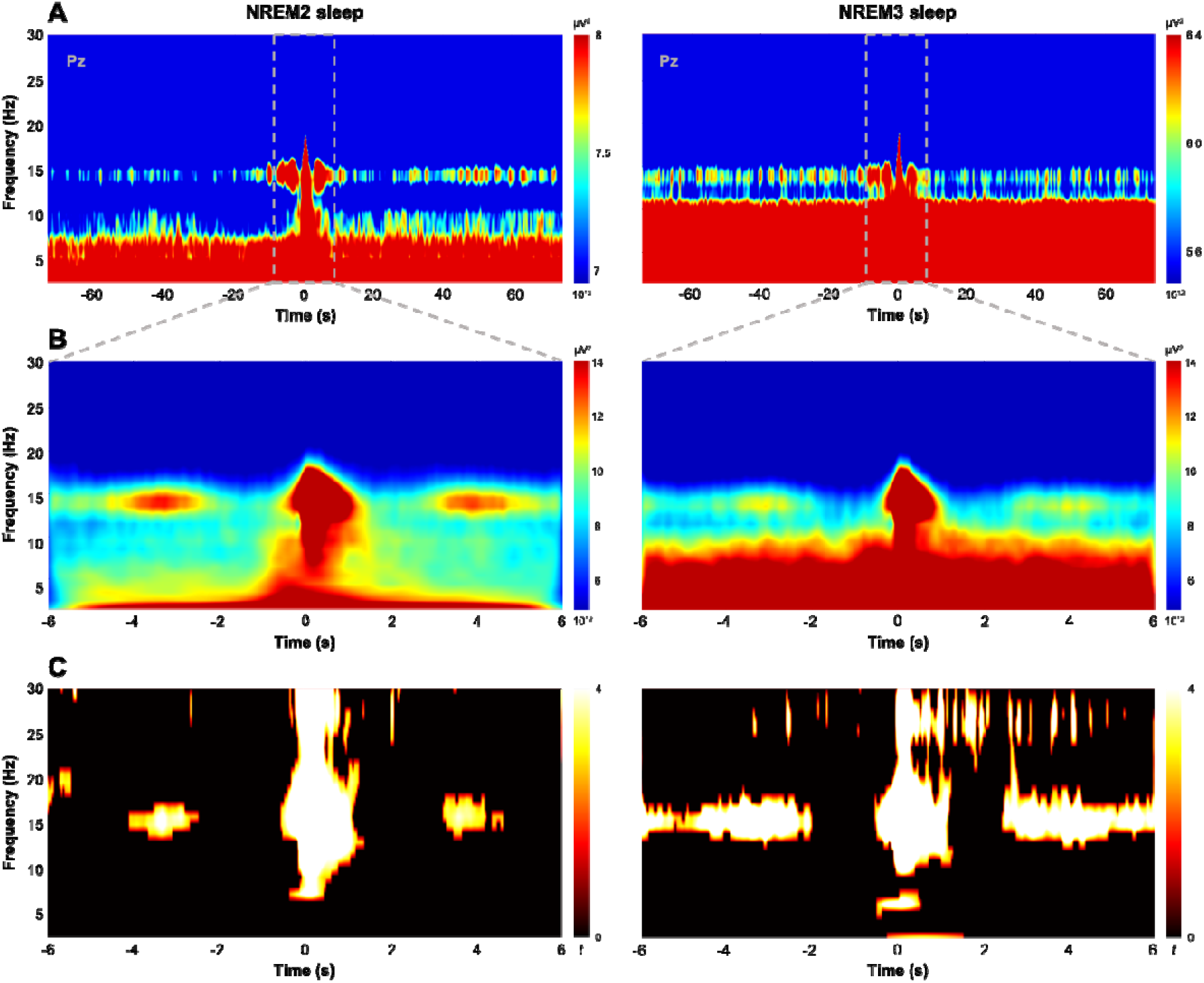
Time-frequency decomposition of NREM2 and NREM3 sleep spindles. (**A**) Grand average time-frequency (TF) maps across participants for spindle events occurring at scalp electrode Pz during NREM2 (left panel) and NREM3 (right panel) sleep periods, using epoch windows ranging from −75 s to +75 s around spindle onsets, and illustrating the time scale of spindle clustering. The color bar reflects spectral power values (_μ_V^2^). (**B**) Grand average TF maps zoomed in on −6 s and +6 s around spindle onsets, illustrating spindle rhythmicity within trains during NREM2 (left panel) and NREM3 (right panel) sleep periods. The color bar reflects spectral power values (_μ_V^2^). (**C**) Statistically significant changes from pre-spindle baseline. The color-coded TF map indicates t-score values and is corrected for multiple comparisons using the Benjamini-Hochberg procedure to control the false discovery rate, *P* < 0.05, N_tests_ = 5581860). NREM: Non-Rapid Eye Movement.

Figure 2B depicts the grand average TF maps zoomed in on −6 s and +6 s around spindle onsets during NREM2 and NREM3 sleep periods. First, TF analysis confirmed the clustering and rhythmic nature of spindle events, with a mean recurrence of 3 ± .4 seconds during NREM2 sleep and 3.2 ± .6 seconds during NREM3 sleep; the ISI difference between NREM2 and NREM3 sleep being non-significant, *t*(33) = −1.26, *p* = .22. The group-averaged TF map illustrates this periodic power increases in the spindle frequency band, which becomes apparent mainly during NREM2 sleep, in comparison to NREM3 sleep due to greater occurrences of spindles in isolation. Nonetheless, statistical TF maps in Figure 2C illustrate the finely tuned and regular occurrence of spindles in trains during both NREM2 and NREM3 sleep, albeit with a more diffuse pattern for the latter.

#### Spindle metrics

To follow-up TF analyses, we correlated spindle density scores (spindle global density [SGD] and spindle local density [SLD] metrics extracted at scalp derivation Pz) with the magnitude of overnight performance gains (see Supplementary Figure 1 for additional analyses across several electrodes of interest). Consistent with recent work, Pearson correlation analyses revealed that SLD correlated positively with the magnitude of performance gains during NREM2 sleep (*r* = .52, *p* = .018) but not during NREM3 sleep (*r* = .43, *p* = .11) (see Figure 3). In contrast, no significant correlations were observed between SGD and performance gains, neither during NREM2 sleep (*r* = .16, *p* = .50) nor during NREM3 sleep (*r* = .38, *p* = .16). Interestingly though, this latter finding suggests that the clustering in time of NREM2 sleep spindles, indirectly reflected by spindle local density, may be of critical importance for efficient consolidation of newly formed motor memories.

**Figure 3.**
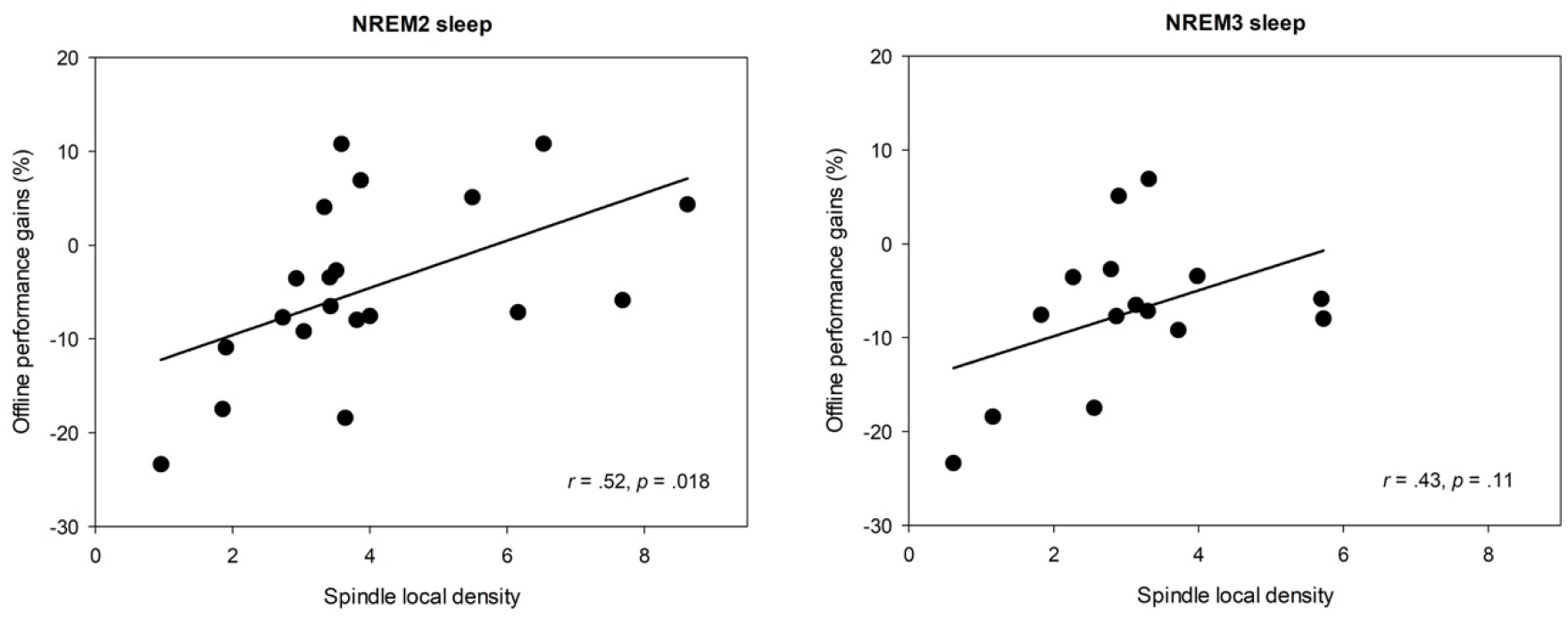
Spindle density and MSL consolidation. Correlations between overnight performance improvements and spindle local density (mean number of spindles within a spindle-centered sliding window of 60 seconds) at the parietal midline derivation Pz during NREM2 sleep (left panel) and NREM3 sleep (right panel). Scatter plots and linear trend lines are provided. Pearson correlation coefficients (*r*) and significance level *p*-values are reported for each correlation. NREM: Non-Rapid Eye Movement.

Hence, adopting a new time-based clustering approach, we determined whether the repetitive and closely spaced in time occurrence of spindles in trains confers favorable conditions for efficient memory reprocessing and consolidation. We thus separately correlated the length of spindle trains and the proportion of grouped over isolated spindles, extracted at scalp derivation Pz, with the magnitude of offline performance gains (see Supplementary Figure 2 for additional analyses across several electrodes of interest). First, analyses revealed that the magnitude of offline performance gains correlated positively with the length of spindle trains during NREM2 sleep (*r* = .61, *p* = .005), but not during NREM3 sleep (*r* = .36, *p* = .19) (see Figure 4); the difference in length of spindle trains during NREM2 sleep (M = 3 ± .7 spindles per train) and NREM3 sleep (M = 2.7 ± .4 spindles per train) being not significant, *t*(33) = 1.31, *p* = .20. Second, significant correlations were observed between offline performance gains and the proportion of grouped spindles during NREM2 sleep (*r* = .67, *p* = .002), while only a trend towards significance was found during NREM3 sleep (*r* = .50, *p* = .06) (see Figure 5); the proportion of grouped spindles differing significantly between NREM2 sleep (M = 60.6 ± 13.5 %) and NREM3 sleep (M = 47.5 ± 14.7 %), *t*(33) = 2.72, *p* = .01, *d* = .93. Thus, altogether our results strongly suggest that the clustering in trains of consecutive spindles during NREM2 sleep may be a critical mechanism underlying the consolidation of motor memories, with spindles occurring in trains playing a more crucial role than isolated ones.

**Figure 4.**
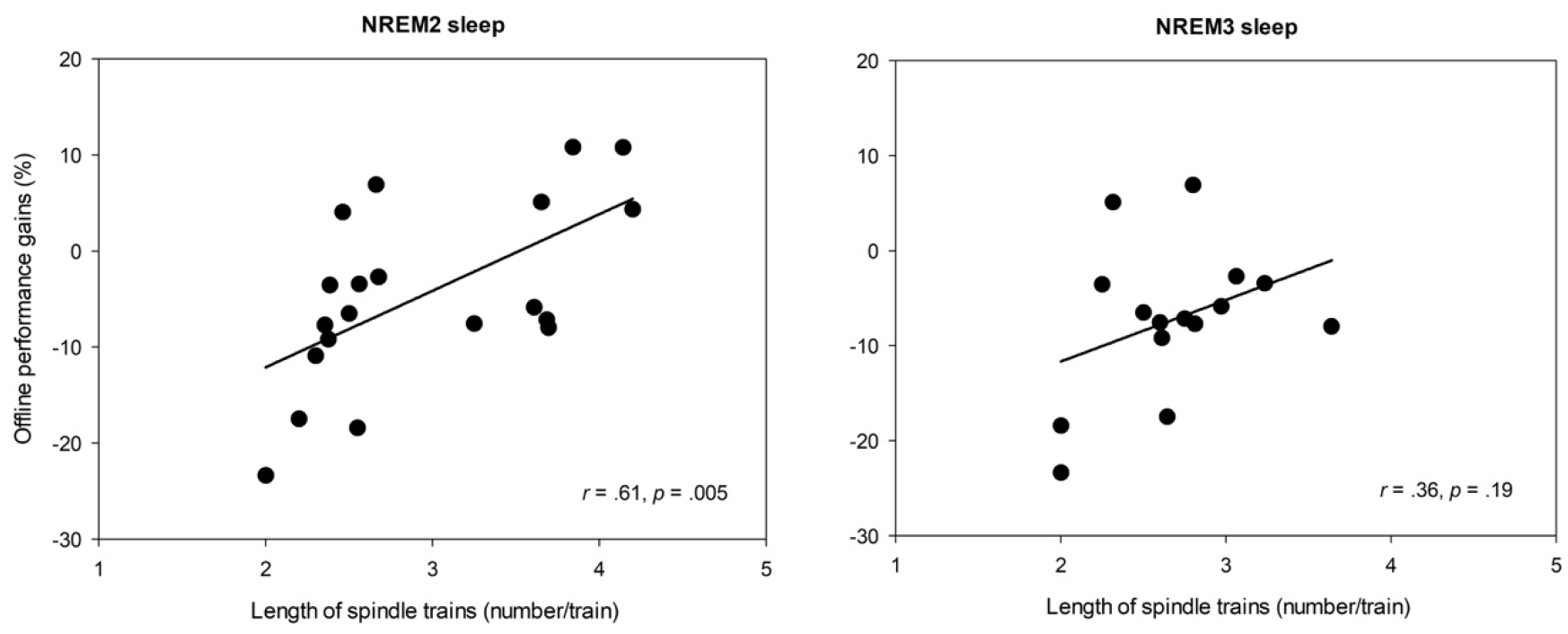
Length of spindle trains and MSL consolidation. Correlations between overnight performance improvements and the length of spindle trains (mean number of spindles per train) at the parietal midline derivation Pz during NREM2 sleep (left panel) and NREM3 sleep (right panel). Scatter plots and linear trend lines are provided. Pearson correlation coefficients (*r*) and significance level p-values are reported for each correlation. NREM: Non-Rapid Eye Movement.

**Figure 5.**
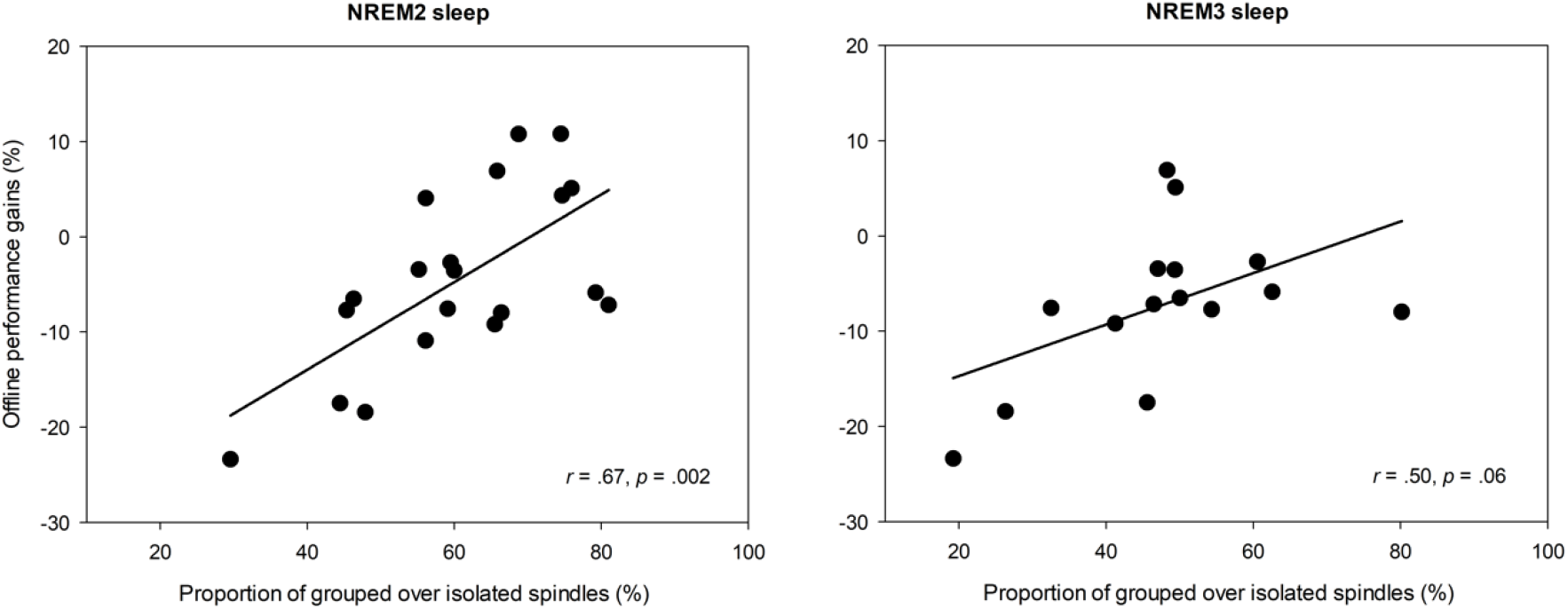
Proportion of grouped over isolated spindles and MSL consolidation. Correlations between overnight performance improvements and the proportion of grouped spindles (in %) at the parietal midline derivation Pz during NREM2 sleep (left panel) and NREM3 sleep (right panel). Scatter plots and linear trend lines are provided. Pearson correlation coefficients (*r*) and significance level p-values are reported for each correlation. NREM: Non-Rapid Eye Movement.

### 2.4. Functional connectivity analysis

Recent work revealed that motor sequence consolidation relies on the synchronized local reprocessing of new information, time-locked to spindles, across segregated but inter-connected brain regions involved in the initial learning process, such as the hippocampus, putamen, thalamus, and motor-related cortical areas^7,8,13^. Here, we wanted to determine whether such network-specific synchronized oscillatory activity during sleep consolidation is related to the rhythmic occurrence of spindles in trains.

#### EEG coherence spindle patterns

We computed coherence (11-16 Hz) patterns for grouped and isolated spindles during NREM2 and NREM3 sleep, and assessed whether they were related to subsequent overnight behavioral changes (see Supplementary Table 1). To that end, we separately regressed iCoh maps of grouped and isolated spindles with the performance gains. Figure 6 shows the connectivity maps of the hippocampus and putamen during NREM2 sleep in relation to the performance gains (Grouped>isolated spindle contrast; see Table 2 for details on significant clusters of activation).

**Table 1.** Experimental sleep characteristics. Sleep architecture during the 2.25-hr sleep interval of the experimental night. Mean, standard errors of the means, minimum and maximum values are reported in minutes. NREM: Non-Rapid Eye Movement, REM: Rapid Eye Movement.

| Sleep stage | Mean | SEM | Min | Max |
| --- | --- | --- | --- | --- |
| Total Sleep Time | 88.30 | 6.25 | 40.5 | 118.50 |
| Awake | 23.63 | 4.15 | 3.5 | 65 |
| NREM1 | 8.60 | 1.46 | 1.5 | 24 |
| NREM2 | 47.50 | 4.45 | 15.5 | 86 |
| NREM3 | 30.88 | 4.90 | 0 | 67 |
| REM | 1.33 | 0.95 | 0 | 18.50 |

**Table 2.** EEG coherence differences between grouped and isolated sleep spindles in relation to MSL consolidation. Statistically significant regressions between hippocampus-seeded and putamen-seeded functional connectivity during grouped and isolated spindles against overnight performance gains (Grouped>Isolated and Isolated>Grouped spindle contrasts). Results are displayed separately for NREM2 and NREM3 sleep.

| Hemisphere | Region | k | x | y | z | T | P (FWE-SVC) |
| --- | --- | --- | --- | --- | --- | --- | --- |
| 1. NREM2 sleep |  |  |  |  |  |  |  |
| 1.1. Regression between functional connectivity of the hippocampus and offline performance gains |  |  |  |  |  |  |  |
| Grouped>Isolated contrast |  |  |  |  |  |  |  |
| Right | Supplementary motor area | 103 | 17 | 5 | 65 | 3.14 | .018 |
|  |  |  | 17 | 3 | 61 | 3.09 | .020 |
| Right | Supramarginal gyrus | 249 | 63 | -21 | 19 | 2.95 | .025 |
| Right | Thalamus | 173 | 5 | -5 | 9 | 2.86 | .029 |
|  |  |  | 9 | -13 | 7 | 2.58 | .047 |
| Left | Thalamus | 17 | -3 | -7 | 11 | 2.62 | .044 |
| Left | Secondary somatosensory cortex/Parietal operculum | 129 | -55 | -21 | 13 | 2.65 | .042 |
|  |  |  | -49 | -19 | 13 | 2.61 | .045 |
| Isolated>Grouped contrast |  |  |  |  |  |  |  |
| No voxels significantly active |  |  |  |  |  |  |  |
| 1.2. Regression between functional connectivity of the putamen and offline performance gains |  |  |  |  |  |  |  |
| Grouped>Isolated contrast |  |  |  |  |  |  |  |
| Right | Thalamus | 198 | 13 | -33 | 1 | 4.14 | .002 |
|  |  |  | 15 | -25 | -1 | 4.13 | .002 |
| Right | Supplementary motor area | 246 | 17 | -3 | 65 | 4.02 | .003 |
| Left | Thalamus | 197 | -9 | -33 | 5 | 3.39 | .010 |
|  |  |  | -5 | -27 | 5 | 3.32 | .011 |
|  |  |  | -7 | -25 | 1 | 3.30 | .012 |
|  |  |  | -15 | -29 | 9 | 3.28 | .012 |
|  |  |  | -15 | -27 | 1 | 3.25 | .013 |
| Right | Hippocampus | 64 | 31 | -25 | -11 | 3.39 | .010 |
|  |  |  | 29 | -33 | -7 | 3.18 | .014 |
| Isolated>Grouped contrast |  |  |  |  |  |  |  |
| No voxels significantly active |  |  |  |  |  |  |  |
| 2. NREM3 sleep |  |  |  |  |  |  |  |
| 2.1. Regression between functional connectivity of the hippocampus and offline performance gains |  |  |  |  |  |  |  |
| Grouped>Isolated contrast |  |  |  |  |  |  |  |
| Right | Supramarginal gyrus | 139 | 61 | -25 | 49 | 2.95 | .024 |
|  |  |  | 61 | -31 | 49 | 2.79 | .030 |
| Isolated>Grouped contrast |  |  |  |  |  |  |  |
| No voxels significantly active |  |  |  |  |  |  |  |
| 2.2. Regression between functional connectivity of the putamen and offline performance gains |  |  |  |  |  |  |  |
| Grouped>Isolated contrast |  |  |  |  |  |  |  |
| No voxels significantly active |  |  |  |  |  |  |  |
| Isolated>Grouped contrast |  |  |  |  |  |  |  |
| No voxels significantly active |  |  |  |  |  |  |  |

Note: k = cluster size (determined at a threshold of $P = .05$ after small-volume correction). Minimum cluster size = 5 voxels. FWE-SVC = family-wise error small-volume correction. x, y, and z coordinates are specified in the MNI space.

**Figure 6.**
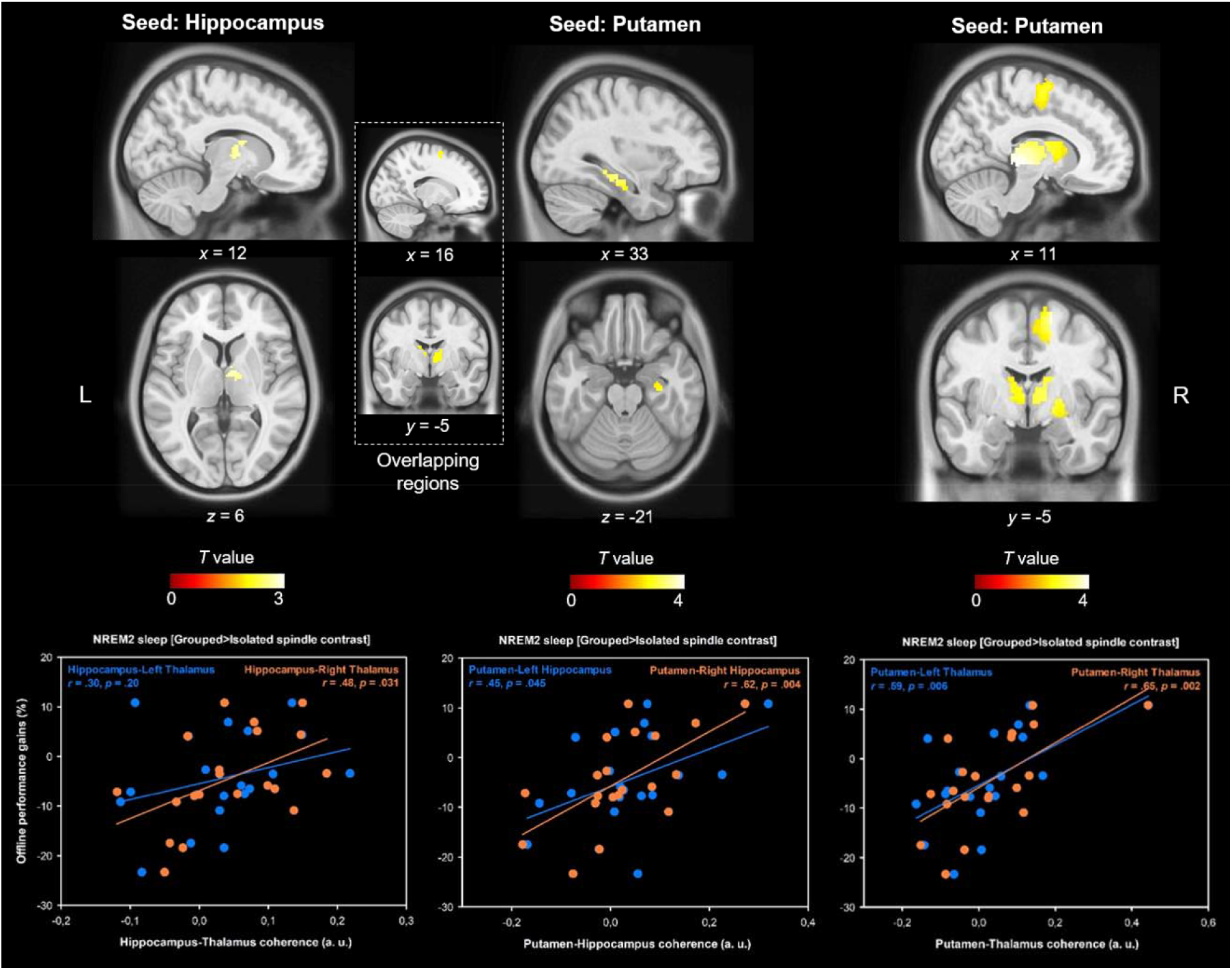
EEG coherence networks during NREM2 sleep spindles. Regression analyses revealed that greater hippocampus- and putamen-seeded coherence in the EEG spindle frequency band (11-16 Hz) during grouped spindles, compared to isolated spindles, was associated with higher overnight performance gains. *Left panel*. As depicted in the scatter plot, hippocampus-seeded connectivity with the right-lateralized thalamus was significantly related to the magnitude of overnight performance gains. *Middle panel*. Putamen-seeded connectivity with the bilateral hippocampus was significantly related to the extent of sleep-dependent performance gains, as depicted in the scatter plot. *Right panel*. Putamen-seeded connectivity with the bilateral thalamus and right-lateralized SMA was significantly related to the magnitude of overnight performance gains, as depicted in the scatter plot. *Top panel*. All images are displayed at one-tailed *p* < .005, uncorrected. Middle-left insets outline the overlapping activations between hippocampus- and putamen-seeded regression maps, which include the bilateral thalamus, caudate and right SMA. See Table 2 for details on significant clusters of activation.

#### Seed: Hippocampus – NREM2 sleep

As expected, regression analyses indicated that increased strength of the functional connectivity (coherence 11-16 Hz) between the hippocampus and several brain regions involved in the MSL consolidation network were linearly related to the offline gains in performance, mainly for grouped spindles. Specifically, for grouped spindles, this brain network consisted of a set of subcortical structures and cortical areas that includes the bilateral thalamus, the SMA, and the parietal cortex. For isolated spindles, only increased coherence between the hippocampus and M1 was significantly and positively related to the offline performance gains. Interestingly though, regression analyses revealed that greater coherence between the hippocampus and the SMA during isolated spindles was negatively associated with offline performance gains, thus suggesting that spindles occurring in isolation may further impair MSL consolidation by way of ineffective memory reactivations.

In addition, we contrasted the iCoh images of grouped and isolated spindles (Grouped>Isolated contrast). We regressed the contrast images against the performance gains to determine whether increases in hippocampus-seeded connectivity during grouped spindles were related to MSL consolidation (Table 2). Contrast results revealed that MSL consolidation was positively related to greater hippocampus-seeded coherence during grouped than isolated spindles, within a brain network including the bilateral thalamus and frontal and parietal cortical regions.

#### Seed: Putamen – NREM2 sleep

Regression analyses indicated that increased strength of the coherence between the putamen and several brain regions involved in the MSL consolidation network was also linearly related to the performance gains for grouped spindles only. Specifically, for grouped spindles, the results demonstrated the involvement of a subcortical-cortical brain network, including the bilateral thalamus, right hippocampus, and parietal regions of the cortex. No significant clusters were found for isolated spindles. However, as for hippocampus-seeded regression analyses, findings indicated that greater coherence between the putamen and the SMA during isolated spindles was negatively associated with MSL consolidation. Finally, contrast results showed that MSL consolidation was positively related to the greater putamen-seeded coherence during grouped than isolated spindles, within a brain network including the bilateral thalamus, right hippocampus, and the right-lateralized SMA (Grouped>Isolated contrast).

Finally, the inset in Figure 6 outlines the overlapping activations between hippocampus-seeded and putamen-seeded contrast regression maps during NREM2 sleep. The overlapping brain regions included the bilateral thalamus, caudate, and right-lateralized SMA. These findings demonstrate the functional contribution of grouped over isolated NREM2 sleep spindles in motor sequence consolidation.

#### Seed: Hippocampus – NREM3 sleep

Regression analyses only revealed that increased strength of the coherence between the hippocampus and the parietal cortex (the supramarginal gyrus) during grouped spindles was positively related to the offline performance gains. No significant cluster of activated voxels was found for isolated spindles. However, as for NREM2 sleep, regression analyses revealed that greater coherence between the hippocampus and MSL-related cortical regions during isolated spindles, including the precuneus and M1, was negatively related to the offline performance gains.

#### Seed: Putamen – NREM3 sleep

Regression analyses failed to detect any significant cluster of activated voxels related to MSL consolidation for both grouped and isolated spindles. Moreover, as for hippocampus-seeded regression analyses, findings indicated that greater coherence between the putamen and M1 during isolated spindles was negatively associated with MSL consolidation.

## 3. Discussion

The aim of the present study was twofold: (i) to determine whether the clustering and multiscale rhythmicity of sleep spindles mediate the consolidation of motor memories, and (ii) to identify the functional contribution of spindles, either occurring in train or in isolation, in the sleep consolidation process. First, our findings corroborated our previous findings^7^, and confirmed the critical role of NREM2 over NREM3 sleep spindle activity in motor memory consolidation. Adopting a new time-based clustering approach, we extended these findings by revealing that spindles occurring in trains during NREM2 sleep play a more critical role than isolated ones in motor memory consolidation. Indeed, through deep brain EEG coherence analyses in the 11-16 Hz frequency range, we revealed that the rhythmic occurrence of spindles in trains might confer optimal neural conditions for efficient reprocessing and consolidation of memory traces within a functional network integrating the hippocampus, the striatum (i.e., the putamen in particular), the thalamus, and motor-related cortical regions. Altogether, our findings suggest that spindles’ clustering and rhythmic occurrence in trains may serve as an intrinsic rhythmic sleep mechanism for the timed reactivation and subsequent consolidation of newly formed motor memories during NREM2 sleep, through synchronized oscillatory activity within a subcortical-cortical network involved during initial learning.

A large body of research has provided substantial evidence that sleep, and spindles in particular, facilitates neuroplasticity and the associated consolidation of both declarative and motor memories^29,40^. Recent empirical research and theoretical frameworks have also outlined the essential contribution of the clustering and hierarchical rhythmicity of sleep spindle activity in the memory consolidation process^13,20,21,23^, especially during NREM2 sleep when it relates to the consolidation of a newly learned motor skill^7,9,19,41^. The present results are thus confirming the positive effects of NREM2 sleep spindles in the consolidation of motor sequences. Also, they accord with the recent theoretical viewpoint outlining the multi-scale periodicity of spindle activity in MSL consolidation. Indeed, time-frequency analyses revealed an apparent clustering and temporal organization of spindle activity during NREM sleep: spindles tend to cluster on a low-frequency time scale of about 50 s (~0.02 Hz rhythm), and during which spindles iterate every 3-4 s (~0.3 Hz rhythm). Yet our findings suggest that such a cluster-based organization and rhythmic occurrence of sleep spindles over MSL-related brain regions is related to memory consolidation only when these occur during NREM2 sleep but not during NREM3 sleep. The latter finding is consistent with a growing body of work proposing that spindles allow for segmenting sleep into clusters of narrow time windows for interspersed and interference-free timed reactivations of memory traces^13,20,29,42^. Consistent with this notion, our results further reveal that the magnitude of performance gains following sleep is positively related to the length of spindle trains during NREM2 sleep, hence leading to the assumption that spindle trains correspond to the underlying neurophysiological mechanisms supporting the repetitive reactivation of previously learned materials^43^. More research is needed to determine the specific content of the memory traces reactivated during spindles occurring in trains, compared to those that arise in isolation.

As aforementioned, we provide here converging evidence in support of the previously reported infra-slow 0.02 Hz periodicity of spindle activity during NREM sleep^20,27^ and extend this finding towards the motor memory domain. Compared to isolated spindle events, it has been suggested that such a temporal clustering of spindles is playing a critical role in the subdivision of NREM sleep into periods of online environmental alertness associated with the gating of sensory information, as well as periods of low fragility to noise crucial for offline memory processing and consolidation^27,29^. Interestingly, however, previous work has demonstrated that such infra-slow oscillatory patterns in cortical EEG rhythms during sleep are coordinated with noradrenergic activity of the locus coeruleus in the brainstem and modulations of the autonomic system, such as cardiac activity and hemodynamic fluctuations^27–29^. Hence, this physiological sleep rhythm may serve a more central regulatory function likely gated by endogenous oscillators involved in the coordination of autonomic outputs with cortical states, possibly through the combined activity of different generators within hypothalamic, subthalamic, and brainstem neural pathways^27,29,44,45^.

In addition to the periodic clustering of spindles in trains, our findings also revealed that spindles tend to recur approximately once every 3-4 seconds during trains. Conceptually, the inter-spindle intervals during trains correspond to periods of refractoriness that are presumably relevant for the timely and orderly segregation of spindle-induced memory reactivations^20^. Indeed, previous studies have shown that the hyperpolarization-activated inward cation current (*I*_h_) plays a key role in the generation and inhibition of rhythmic activities in thalamocortical cells^46,47^. The deactivation of *I*_h_ is believed to cause the fading of spindles and to prevent further thalamocortical oscillations within defined refractory periods. Hence, in accordance with previous work, our findings reveal that spindle recurrence is limited in time over intervals of about 3-4 seconds during NREM sleep, interspaced by refractory periods. They further support the notion that such a ~0.3 Hz rhythm of spindle incidence during trains is likely supported by a combination of intra-thalamic, cortical, and brainstem mechanisms (see ^13,23,29^ for reviews).

More importantly, our current results strongly suggest that sleep mechanisms underlying the topographical expression of spindles within clustered and temporally organized patterns during NREM2 sleep may be fundamental for the local reprocessing and subsequent consolidation of interrelated memory units into broader, coherent representations (see also Supplementary Figures 1–2). Although the physiologic origins of this timed occurrence of spindles in trains are not fully understood and thus require further research, our findings point towards a physiological advantage of spindle clustering and rhythmicity in the sleep-dependent consolidation of motor memories.

Our findings are congruent with the theoretical assumption that sleep spindles occurring in trains may play a more critical role than isolated ones in motor sequence consolidation^13^. Here, we directly tested this hypothesis by determining the functional contribution of grouped and isolated spindles in the consolidation process. Through deep brain EEG coherence analyses, time-locked to spindles, our findings reveal that the magnitude of overnight performance gains is related to the repeated reactivation of functionally related brain structures within a subcortical-cortical network involved in motor sequence learning, such as the hippocampus, the striatum (the putamen in particular), the thalamus, and motor-related cortical circuitry. Hence, our findings corroborate previous research revealing that oscillatory synchrony in the EEG spindle frequency band involves cross-structural reactivation and communication between MSL-specific brain regions during sleep consolidation^7,8,11^. In addition, we further demonstrate that this MSL-related strengthened functional network is related to the rhythmic occurrence of spindles in trains during NREM2 sleep, thus revealing that grouped spindles play a more functional role than isolated ones in motor memory consolidation. From a holistic standpoint, we therefore advocate for a broader and multi-dimensional approach of sleep spindle activity in future research on memory consolidation, taking into account the hierarchical clustering and multiscale rhythmicity of sleep spindles rather than their mere overall occurrence, as well as other characteristics such as amplitude, duration, and frequency metrics.

Our results complement the extant literature on motor sequence learning and sleep-related consolidation in several ways^8,11,17,34^. Previous studies have revealed that NREM2 sleep spindles may play a causal role in motor memory consolidation^19,48^, possibly through their ability to induce synaptic potentiation and plasticity^49,50^ via reactivation of the memory trace^8^. Here, we provide further evidence that not all spindles contribute to the consolidation process. On the contrary, our results suggest that sleep spindles occurring in trains are more important in enhancing memory consolidation by locally and repetitively reactivating task-relevant brain regions that were engaged during the learning process *per se*. This finding is consistent with the “synaptic consolidation” hypothesis, which refers to transforming previously learned information into a long-term form at the local synaptic level^6,51^. In addition, our results add to a growing body of work linking the interaction between the striatal and hippocampal memory systems during sleep to the consolidation of motor memories^34,52,53^. More specifically, we assume that the repetitive engagement and interaction of the cortico-striatal and cortico-hippocampal circuits during spindle trains subserves sleep-dependent MSL consolidation. This assumption further agrees with recent studies showing a gradual reorganization of motor representations at the system level towards a subcortically-dominant consolidated memory trace during NREM sleep^11,17^; with synaptic consolidation mechanisms serving as subroutines for the systems-level reorganization of memory traces^51,54^. Hence, we conjecture that the clustering of spindles in trains during NREM2 sleep would serve as an intrinsic rhythmic sleep mechanism allowing the repeated global synchronization and communication within cortical and subcortical networks, thus supporting long-range synaptic consolidation and memory transfer to distant cortical and subcortical regions^11,55,56^.

Yet, in contrast to the findings observed for grouped spindles, regression analyses revealed significant negative relationships between performance gains and coherences of the hippocampus and the putamen with MSL-related cortical regions (mainly right SMA and left M1) during isolated spindles. These latter results suggest that spindles occurring in isolation during NREM sleep may rather impair MSL consolidation through ineffective, non-recurring memory reactivations. Hence, while spindles occurring in trains can strengthen memory representations through timed reactivations, in contrast, spindles occurring in isolation may instead activate sleep mechanisms underlying the clearance or decreased accessibility of the memory content (see ^57^ for a review). One potential explanation would be that isolated spindles may serve another memory processing function, such as forgetting^58–61^. Sleep spindles may weaken memory representations when occurring in isolation through less frequent reactivations during periods where depressive plasticity is favored, thus instigating forgetting through the erasure of region-specific or interrelated memory units. Further research is therefore needed to unravel the physiologic origins and memory functions of grouped and isolated spindles, and the extent to which brain (re)activations may favor or impair motor memory consolidation.

To conclude, our findings confirm the critical role of NREM2 over NREM3 sleep spindle activity in motor memory consolidation. Interestingly, our results point out the role of the temporal cluster-based organization of NREM2 sleep spindles in the consolidation process, as sleep spindles tend to cluster on a low-frequency time scale of about 0.02 Hz, with spindles iterating at an intermediate rhythm of about 0.3 Hz during spindle trains. In addition, the current results suggest that the rhythmic occurrence of spindles in trains serves as an intrinsic clocking sleep consolidation mechanism underlying the timed and repeated reactivation of functionally related brain structures within a subcortical-cortical network involved in motor sequence learning, such as the hippocampus, the striatum (the putamen in particular), the thalamus, and motor-related cortical circuitry.

## 4. Materials and methods

### 4.1. Ethics Statement

The study protocol was approved by the Research Ethics Board of the “Regroupement Neuro-imagerie Québec” (CMER RNQ 13-14-011). Before participation, all participants provided written informed consent and received financial compensation for participating in this study.

### 4.2. Participants and procedure

Thirty-three young healthy volunteers (mean age: 25.5 ± 3.7 years, 24 females) were recruited through local advertisements. All participants included in the study fulfilled the following inclusion criteria: right-hand dominant (tested using the Edinburgh Handedness Inventory questionnaire^62^), non-smoker, non-musician or typist, medication-free, no history of mental illness, no signs of mood disorders (i.e., self-rated score ≤ 8 on the Beck Depression^63^ and Anxiety Inventories^64^), Body Mass Index ≤ 25, and no sleep disturbances (i.e., Pittsburgh Sleep Quality Index Global Score < 5^65^). Participants categorized as extreme morning or evening types (tested with the Horne Ostberg Morningness-Eveningness Scale^66^), shift workers, or those who experienced a trans-meridian trip less than three months before the experiment were excluded. Furthermore, we also excluded participants exhibiting signs of excessive daytime sleepiness (≤ 9 on the Epworth Sleepiness Scale^67^). Participants were asked to maintain a regular sleep-wake cycle (bedtime between 10:00 pm–12:00 am, wake-time between 07:00 am–10:00 am) to ensure they were not sleep-deprived. The schedule’s compliance was assessed using sleep logs and wrist-worn actigraphs (Actiwatch 2, Philips Respironics, Andover, MA, USA). Finally, volunteers were also asked to refrain from all caffeine- and alcohol-containing beverages 24 hours before and during the experimental period.

Each participant completed three visits to the Functional Neuroimaging Unit (UNF) located at the Centre de Recherche de l’Institut Universitaire de Gériatrie de Montréal (CRIUGM). Visit 1 served as a polysomnographic (PSG) screening and acclimatization night, whereas visit 2 (training+sleep) and visit 3 (retention) consisted of the experimental sessions. Visits 1 and 2 were separated by approximately seven days, while visits 2 and 3 were administered twenty-four hours apart. During visit 1, participants were given a 2.25-hour opportunity to sleep in a mock scanner. The noise and lighting conditions were almost identical to that experienced in the actual 3.0T TIM TRIO MRI scanner (Siemens, Erlangen, Germany) located at the UNF, CRIUGM. During the screening and experimental nights, simultaneous EEG-fMRI data were recorded using an MR-compatible EEG cap (EasyCap, Brain Products). Based on screening PSG data, participants were included in the study only if they met the arbitrary criteria of 15 min of consolidated NREM2 sleep (considered the minimum amount of data necessary for data analysis purposes). All screening and experimental EEG sleep recordings started between 11:00 pm and midnight. Sleep recordings ended after a maximum of 2 hours and 15 minutes in both the mock and actual MRI scanners (equivalent to 4000 volumes, corresponding to the maximum number of possible volumes determined by the system for a single fMRI session). Following the sleep-EEG recordings in the mock (visit 1) or actual MR scanners (visit 2), EEG electrodes were removed and participants were allowed to sleep for the rest of the night in the nearby sleep laboratory until 7:00-8:00 am. Of the twenty-six participants who met the 15 min criteria of NREM2 sleep during Visit 1 (mean age: 25.4 ± 3.6 years, 20 females), six were excluded from the data analyses. Three participants had disturbed sleep during the experimental night (i.e., sleep interspersed with periods of wakefulness), and three had low-quality MR-denoised EEG signals for accurate sleep scoring and spindle analysis. The remaining 20 participants (mean age: 25.3 ± 3.5 years, 16 females) met all the inclusion criteria throughout the study, and their data were therefore considered for later data analyses. The present study’s sample size and inclusion rate align with previous EEG-fMRI sleep studies (see ^7,68–70^).

### 4.3. Experimental design and tasks

Using a within-subjects design, overnight motor skill consolidation was assessed by computing the change (%) in performance between two delayed retention tests, performed 15 min and 24 hours after the end of practice. During training and retention sessions, the MSL task was performed while participants were lying supine in the MRI scanner. The participant’s performance on the MSL task was recorded using a custom-made MR-compatible response box composed of four equidistantly spaced and ergonomically located response buttons. All aspects of the experiment were programmed with MATLAB***®*** (The MathWorks, Inc., Natick, MA) using the Psychophysics Toolbox extensions.

#### Motor Sequence Learning task

The MSL task required participants to perform, as rapidly and accurately as possible, a 5-element finger movement sequence by pressing the corresponding buttons on the response pad with fingers of their non-dominant left hand (see ^7,8^ for similar procedures). Response times (RTs) were measured as the interval between consecutive self-initiated (i.e., non-cued) key presses. The sequence to perform (1-4-2-3-1, where 1 corresponds to the little finger and 4 to the index finger) was explicitly taught to the participants before training. Specifically, the session included a brief pre-training phase during which participants were not scanned and instructed to repeatedly and slowly perform the 5-element sequence until they accurately reproduced it three consecutive times. This pre-training phase was intended to ensure that participants understood the instructions and were able to perform the task after they explicitly memorized the sequence.

The training session consisted of fourteen blocks of practice (indicated by a green cross displayed in the center of the screen), each composed of twelve repetitions of the motor sequence (i.e., 60 key presses per block). Practice blocks were interspersed with 25s rest periods (indicated by the onset of a red cross on the screen) to prevent fatigue. The participants were given no information or feedback regarding their performance throughout the training session. First, participants underwent the training session about 30 minutes before going to sleep. They were then administered two delayed retention tests, performed 15 minutes (test) and 24 hours (retest) after the end of training. Each of the two retention tests consisted of only one practice block to minimize the effects of additional practice on the MSL task.

To reflect individual skill levels, we quantified MSL performance with a score derived from a linear speed-accuracy function (see also ^7^). This measure, which presents in a single score a measure sensitive to both speed and accuracy, was calculated by dividing the number of accurately typed sequences by the RT. Performance score (PS) values for individual RTs were then averaged to yield a global estimate of the PS for each practice block. Hence, mean PS was computed for each practice block during the training and testing sessions (see Figure 1). Motor skill acquisition was evaluated by analyzing the within-session performance gains from the first (Block 1) to the last training block (Block 14). Motor skill consolidation was assessed by analyzing the between-session performance gains from the 15-min test to the 24-hr retest.

##### EEG data acquisition and pre-processing

EEG was acquired using a 64-channel MR-compatible EEG cap (including one electrocardiogram [ECG] electrode) (BrainCap MR, Brain Products Inc.). The EEG cap included 63 scalp flat ring-type electrodes (5 kΩ safety resistors), with FCz and AFz being the reference and ground electrodes. In addition to the Braincap ECG electrode (10 kΩ safety resistor; placed left-sided along the paravertebral line between the 5th and 7th costa), bipolar ECG recordings taken from V2-V5 and V3-V6 were acquired via MR-compatible Ag-AgCl electrodes with 10 kΩ safety resistors. EEG and ECG data were recorded using two battery-powered MR-compatible 32-channel amplifiers and a 16-channel bipolar amplifier (respectively, BrainAmp MR and BrainAmp ExG MR, Brain Products Inc.). EEG and ECG signals were digitized at a 5-kHz sampling rate with a 500-nV resolution. Data were transferred outside the scanner room through fiber-optic cables to a personal computer where the EEG system running the Vision Recorder software (v1.03; Brain Products) was phase-synchronized to the MR scanner clock (SyncBox, Brain Products Inc.). Sleep EEG was monitored online using the RecView software (Brain Products). Electrode-skin impedance was kept below 5 kΩ using Abralyt HiCl electrode paste (Easycap) to ensure stable recordings throughout the night. MRI foam cushions were used to fix the participant’s head in the head coil to minimize movement-related EEG artifacts during the night.

EEG data were corrected for MR gradient artifacts using the “fMRI Artifact Slice Template Removal” (FASTR^71^) provided by the fMRIB plug-in for EEGLAB. Cardioballistic pulse-related artifacts were separated from the signal using an independent component analysis approach, constrained by the detected QRS peaks (using the ECG channel with the smallest number of detected QRS outliers), and then removed using the “fMRI Artifact rejection and Sleep Scoring Toolbox” (FASST^72^) implemented in Matlab®. The MR-denoised EEG signal was bandpass filtered between 0.5 and 30 Hz to remove low-frequency drift and high-frequency noise, down-sampled to 250 Hz, and re-referenced to the linked mastoids (i.e., TP9 and TP10).

### 4.4. Brain imaging: MR pulse sequence parameters

Functional and structural images were acquired on a Siemens TIM Trio 3T scanner with two different setups. High-resolution structural T1-weighted MR images were acquired using a 32-channel head coil and a Multi-Echo MPRAGE sequence (MEMPRAGE; voxel size = 1 mm isometric; TR = 2530 ms; TE = 1.64, 3.6, 5.36, 7.22 ms; FA=7°; GRAPPA = 2; FoV = 256 × 256 × 176 mm), with the different echoes combined using a Root-Mean-Square.

Functional MR images were acquired with a 12-channel head coil, whose shape and inner size was more appropriate for sleep and EEG settings. We used an EPI sequence providing complete cortical and cerebellum coverage (40 axial slices, acquired in ascending order, TR = 2160 ms; FoV = 220 × 220 × 132 mm, voxel size = 3.44 × 3.44 × 3.3 mm, TE = 30 ms, FA = 90°, GRAPPA = 2). Following each session of fMRI data acquisition, four volumes were acquired using the same EPI sequence but with a reversed-phase encoding to enable retrospective correction of distortions induced by B0 field inhomogeneity. Also, note that the EPI sequence parameters were chosen so that the gradient artifact would be stable in time and to ensure that the lowest harmonic of the gradient-induced artifact (18.52 Hz) would occur beyond the 11-16 Hz spindle frequency range (see ^7,8^ for methodological details).

### 4.5. PSG recordings and spindle detection

The artifact-free EEG signal was first sleep stage scored according to AASM guidelines^73^. Every 30s epoch of the EEG data was visually scored by a registered polysomnographic technologist as either NREM stages 1-3, REM, or wake (see Table 1 for details). Spindle detection was then conducted using all artifact-free NREM sleep epochs. We restricted the detection of spindle events over the parietal site (Pz electrode), as expression of MSL-related spindles has been shown to predominate over this region^7,19,74^. Discrete sleep spindle events (i.e., onset and offset) were automatically detected using a wavelet-based algorithm (https://github.com/labdoyon/spindlesDetection; see ^7^ for further details). Spindles were detected at the Pz derivation by applying a dynamic thresholding algorithm (based on the standard deviation above the mean in the power spectrum) to the extracted wavelet scale corresponding to the 11-16 Hz frequency range and a minimum window duration set at 300 ms^7,75^. Thus, detected events were considered sleep spindles only if the events lasted 0.3-2 seconds, occurred within the 11-16 Hz frequency range, and with an onset occurring within NREM2 or NREM3 sleep periods. Note that the limited amount of NREM2 and NREM3 sleep periods prevented further subdivision into slow (e.g., ~11-13.5 Hz) and fast (e.g., ~13.5-16 Hz) spindles due to the insufficient number of events per spindle category for statistical relevance (see also ^7^).

### 4.6. Spindle characteristics

For each spindle event, several variables of interest were considered: two measures of spindle density (global density [number of spindles per minute] and local density [number of spindles within a spindle-centered sliding window of 60 seconds]), three metrics related to spindle clustering (number of spindle trains, length of spindle trains [mean number of spindles per train], and the ratio between grouped and isolated spindles [in percentage]), and two measures of spindle rhythmicity (inter-spindle interval [ISI] and inter-train interval [ITI]; in second). Such metrics sensitive to spindle clustering and rhythmicity would reveal whether spindles occur within timely organized patterns to efficiently consolidate specific memories or erratically within irregular patterns across NREM sleep episodes.

Adopting a new time-based clustering approach^13^, we determined whether the repetitive and closely spaced in time occurrence of spindles in trains over specific brain regions may confer favorable conditions for efficient reprocessing and consolidation of memory traces. To that end, and as recently proposed by Boutin and Doyon^13^, trains of spindles were operationalized as clusters of two or more consecutive and electrode-specific spindle events interspaced by less than or equal to 6 s, in comparison to those occurring in isolation (i.e., more than 6 s between two consecutive spindles detected on the same electrode). For convenience, spindles belonging to trains were labeled grouped spindles, and those occurring in isolation were labeled isolated spindles. Thus, two metrics were derived from this time-based clustering: the length of spindle trains [mean number of spindles per train] and the proportion of grouped over isolated spindles [in percent]).

### 4.7. EEG analysis

#### Time-Frequency analysis

To illustrate spindle clustering and rhythmic regularities across NREM sleep periods, time-frequency (TF) analyses of sleep spindles were conducted separately for NREM2 and NREM3 sleep epochs. TF decomposition was done using Morlet wavelet transforms across the entire 1-30 Hz frequency range (μV^2^/Hz) and for epoch windows of ± 75 s around spindle onsets to highlight the time scale of spindle clustering (inter-train interval), and of ± 6 s around spindle onsets to study the rhythmicity of spindles within trains (inter-spindle interval).

#### Source reconstruction

Source localization of spindle-related cortical and subcortical activity was estimated using a deep brain activity (DBA) model, which realistically distributes the current dipoles over the neocortex and subcortical structures. For each participant, head structures (cortex, skull, head) were automatically segmented from T1-anatomical MR images using the FreeSurfer 5.3 software. For subject co-registration and further group analysis, we created a template source grid in the normalized MNI (Montreal Neurological Institute) space with a 4-mm resolution. We projected the grid to the individual subject spaces, yielding subject-specific grids that map onto the template grid in spatially normalized space. Hence, after decimation and down-sampling, individual volume head models were decomposed into ~30 000 voxels (cortex and deep structures), which represented the current dipoles (unconstrained orientations) where the source signal was to be estimated (in pico ampere meter; pA.m). Cortical and subcortical structures known to be involved in either the sleep spindle or MSL-consolidation networks, such as the bilateral thalamus, hippocampus, caudate nucleus, putamen as well as parietal and motor-related cortical regions^4,7,35,68,76^ were included in the volume head model. Using this anatomical and electrophysiological DBA model, the forward model was computed using the OpenMEEG software^77,78^. To estimate the current distributions, we computed the noise-normalized (noise covariance matrix) depth-weighted minimum norm estimate using the Brainstorm software^79^.

#### Functional connectivity analysis

As previously used by Boutin and colleagues^7^, coherence-based metrics were adopted to estimate functional connectivity from EEG data. EEG coherency is a measure that reflects the coupling of frequency spectra between two signals from different brain sites, as defined by their normalized cross-spectral density and relative phase delay. However, to avoid spurious results caused by volume conduction artifacts or signal leakage effects, we computed the imaginary part of coherency (see ^80^), which is considered to reflect true brain interactions by being artifact-free and insensitive to instantaneous synchronized activity (i.e., with no phase difference). Thus, we computed whole-brain source connectivity analysis by estimating the imaginary part of coherency between specific seed regions (e.g., bilateral putamen and hippocampus) and other cortical/subcortical regions for each spindle epoch (between 0 sec and 2 sec). Seed regions were selected *a priori* with reference to the MSL literature and based on recently published EEG-fMRI studies^7,8,10,11^. Individual iCoh maps (source-coherence matrices) were baseline-normalized (from −2000 to −500 ms before spindle onset) and averaged for each participant in the spindle frequency band (11-16 Hz). Before performing group-level analysis, iCoh maps computed on individual brains were back-projected into the MNI stereotaxic space (template source grid).

#### Regression model and iCoh maps

To assess the relationship between spindle-related brain activation and the expression of overnight performance gains from the 15-min test to the 24-hr retest, we regressed individual iCoh maps with their respective offline performance gains. Based on our *a priori* hypothesis and recent studies, regions of interest (ROI) were restricted to MSL-related cortical areas and subcortical structures. Hence, the thalamus, putamen, hippocampus, and caudate nucleus were considered subcortical structures of interest. Relevant cortical ROIs for subsequent analyses were restricted to parietal, pre-motor, and sensorimotor brain areas (the primary motor (M1) and somatosensory (S1) cortices). In addition, to test for the specificity of the selected ROIs in MSL consolidation, iCoh values from subcortical and cortical structures not related to the MSL network (i.e., the bilateral amygdala and caudal middle frontal gyrus, respectively) were also extracted.

## Code availability

Sleep EEG data were processed using the MATLAB R2019b software from The MathWorks (Natick, MA) and the open-source Brainstorm and EEGLAB softwares. The codes for the detection and clustering of sleep spindles are available at the following GitHub repositories: https://github.com/labdoyon/spindlesDetection and https://github.com/arnaudboutin/Spindle-clustering. The codes used to perform other analyses are available from the corresponding author upon reasonable request.

## Data availability

The data supporting the results of this study are available from the corresponding author upon reasonable request.

## Supplementary Notes and Figures

### Spindle density

To determine whether similar spindle density patterns were observed over cortical regions involved in MSL consolidation, we extended correlation analyses of both spindle local density (SLD) and spindle global density (SGD) metrics across several electrodes of interest (F3, F4, Fz, C3, C4, Cz, P3, P4, Pz, O1, O2, Oz). The correlation matrix revealed significant correlations between SLD and performance gains over pre-motor, sensorimotor and parietal cortices (Supplementary Figure 1 shows the correlation matrix heatmaps for both spindle density metrics during NREM2 sleep, respectively the SLD and SGD; correlograms thresholded at p < .05, uncorrected for multiple comparisons). This finding suggests that the repetitive and closely spaced in time occurrence of spindles over brain regions involved during initial learning may relate to the iterative reprocessing of interrelated memory units during sleep-dependent consolidation.

### Spindle clustering

To provide a topographical overview of spindle clustering over the cortex, we extended correlation analyses of both the length of spindle trains and the proportion of grouped over isolated spindles across several electrodes of interest (F3, F4, Fz, C3, C4, Cz, P3, P4, Pz, O1, O2, Oz). The correlation matrices revealed significant correlations between both clustering metrics and the offline performance gains over pre-motor, sensorimotor, and parietal cortices (Supplementary Figure 2 shows the correlation matrix heatmaps for both clustering metrics during NREM2 sleep, respectively the length of spindle trains and the proportion of grouped over isolated spindles; correlograms thresholded at p < .05, uncorrected for multiple comparisons). These results suggest that the sleep mechanisms underlying brain-wide spatial distribution of spindles within clustered and temporally organized patterns during NREM2 sleep may be fundamental for effective local reactivation of learning-related memory units and subsequent consolidation of broader, newly-formed motor memories.

---

**Supplementary Figure 1.**
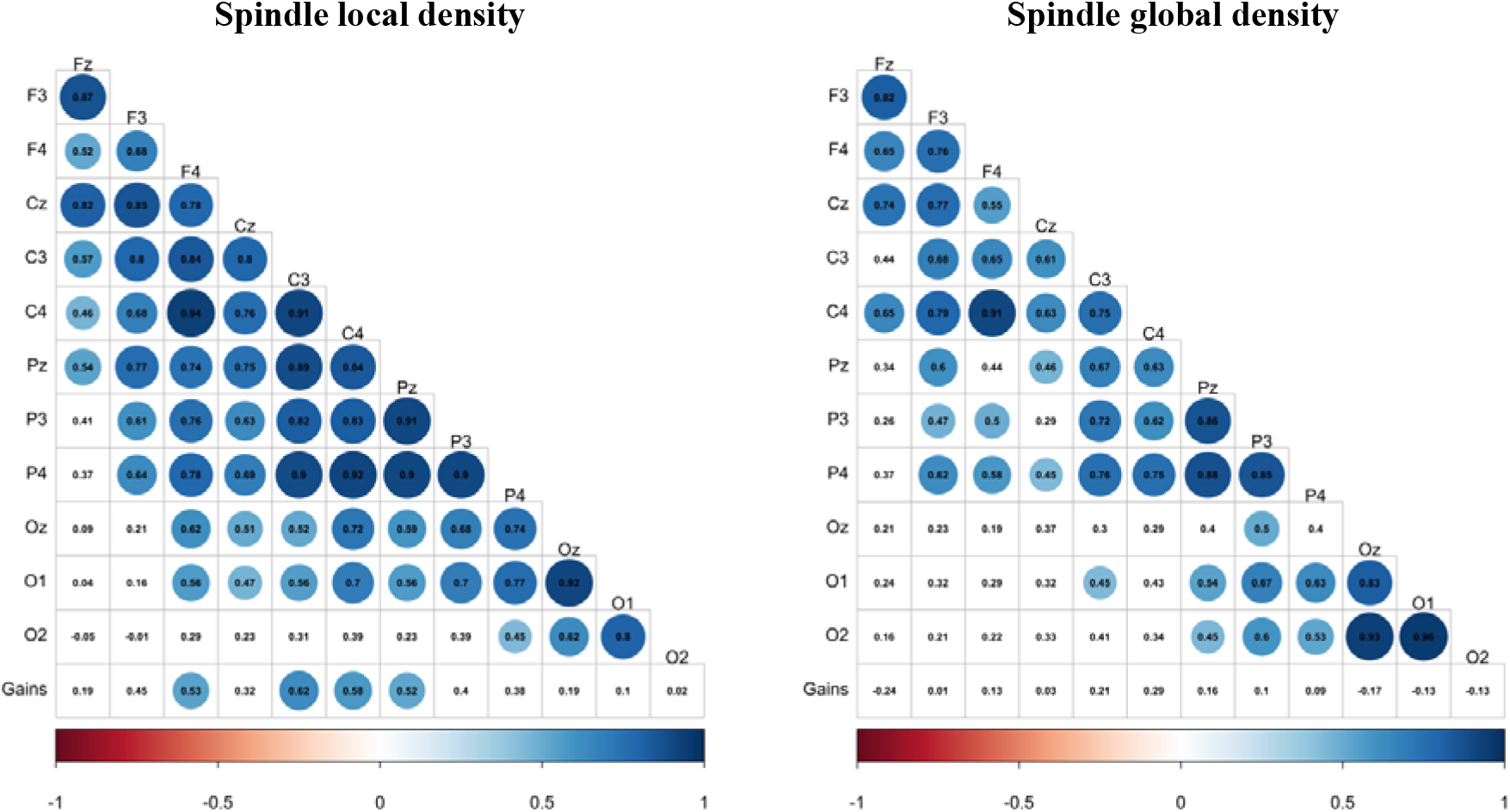
Spindle density and MSL consolidation. Correlations between overnight performance improvements with spindle local density (mean number of spindles within a spindle-centered sliding window of 60 s; left panel) and spindle global density (number of spindles per minute; right panel) detected over main scalp derivations during NREM2 sleep. Pearson correlation coefficients (*r*) and significance level p-values are reported for each correlation. Colored circles highlight significant p-values (*p* < .05, uncorrected for multiple comparisons).

**Supplementary Figure 2.**
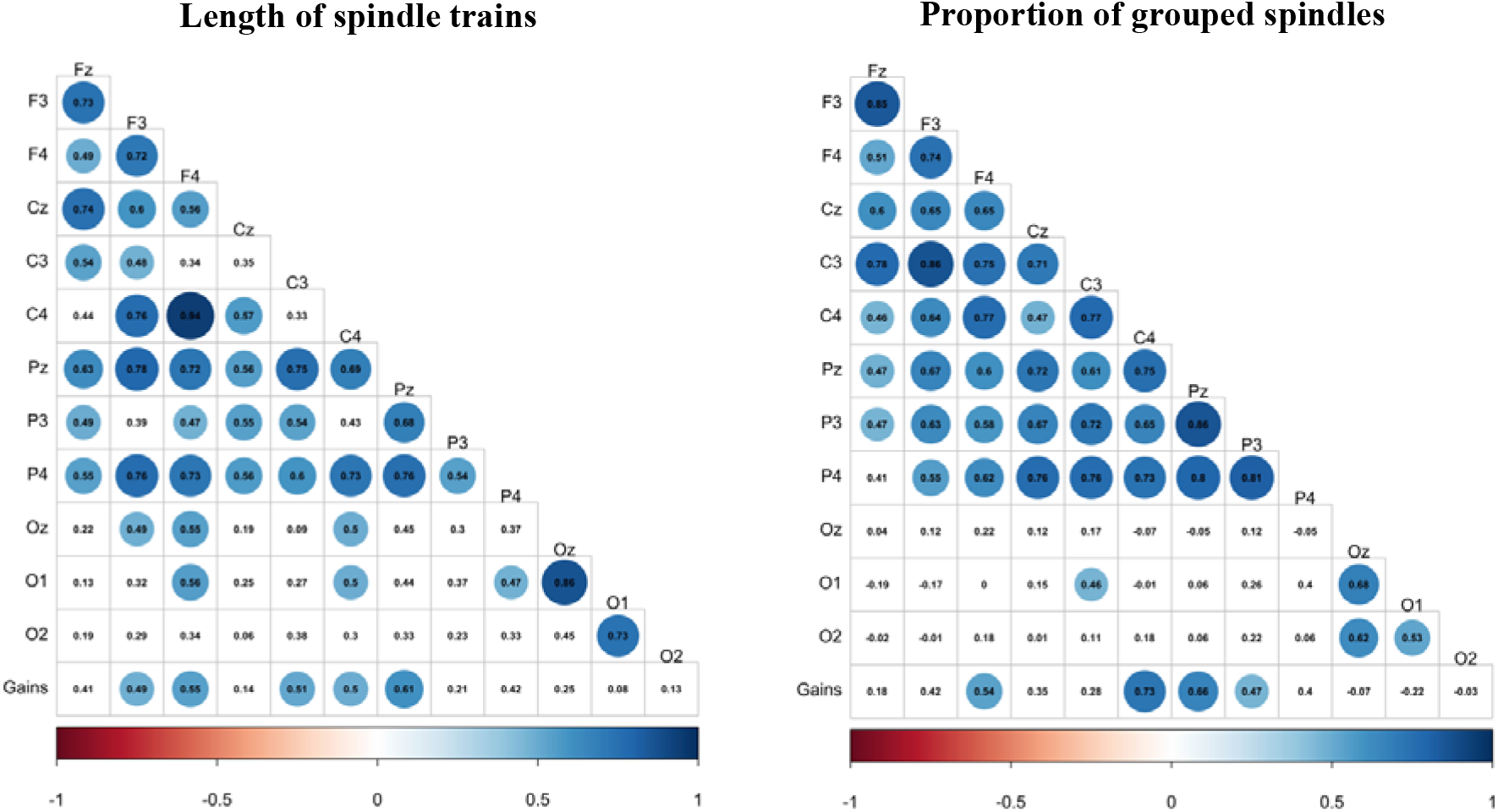
Spindle clustering and MSL consolidation. Correlations between overnight performance improvements with the length of spindle trains (mean number of spindles per train; left panel) and the proportion of grouped spindles (in %; right panel) detected over main scalp derivations during NREM2 sleep. Pearson correlation coefficients (*r*) and significance level p-values are reported for each correlation. Colored circles highlight significant p-values (p < .05, uncorrected for multiple comparisons).

**Supplementary Table 1.** EEG coherence patterns during grouped and isolated sleep spindles in relation to MSL consolidation. Statistically significant regressions between functional connectivity of the hippocampus and putamen during grouped and isolated spindles against overnight performance gains. Results are displayed separately for NREM2 and NREM3 sleep.

| Hemisphere | Region | k | x | y | z | T | P (FWE-SVC) |
| --- | --- | --- | --- | --- | --- | --- | --- |
| **1. NREM2 sleep** |  |  |  |  |  |  |  |
| **1.1. Regression between functional connectivity of the hippocampus and offline performance gains** |  |  |  |  |  |  |  |
| **1.1.1. Grouped spindles** |  |  |  |  |  |  |  |
| **Positive correlation** |  |  |  |  |  |  |  |
| Left | Precuneus | 197 | -7 | -51 | 29 | 3.09 | .020 |
|  |  |  | -3 | -51 | 31 | 3.09 | .021 |
| Right | Thalamus | 154 | 7 | -17 | 17 | 3.01 | .024 |
|  |  |  | 9 | -15 | 13 | 2.98 | .025 |
|  |  |  | 9 | -11 | 15 | 2.91 | .028 |
|  |  |  | 5 | -9 | 13 | 2.90 | .029 |
| Left | Supplementary motor area | 143 | -9 | -13 | 47 | 2.84 | .032 |
| Left | Thalamus | 43 | -13 | -13 | 17 | 2.70 | .040 |
| **Negative correlation** |  |  |  |  |  |  |  |
| No voxels significantly active |  |  |  |  |  |  |  |
| **1.1.2. Isolated spindles** |  |  |  |  |  |  |  |
| **Positive correlation** |  |  |  |  |  |  |  |
| Right | Primary motor cortex | 135 | 35 | -23 | 69 | 2.63 | .040 |
| **Negative correlation** |  |  |  |  |  |  |  |
| Right | Supplementary motor area | 114 | 17 | 5 | 65 | 3.47 | .009 |
| **1.2. Regression between functional connectivity of the putamen and offline performance gains** |  |  |  |  |  |  |  |
| **1.2.1. Grouped spindles** |  |  |  |  |  |  |  |
| **Positive correlation** |  |  |  |  |  |  |  |
| Right | Thalamus | 20 | 23 | -33 | -1 | 3.11 | .015 |
| Right | Hippocampus | 48 | 29 | -33 | -7 | 2.95 | .020 |
|  |  |  | 33 | -31 | -11 | 2.94 | .020 |
|  |  |  | 29 | -39 | -1 | 2.84 | .024 |
| Left | Precuneus | 203 | -9 | -79 | 51 | 2.65 | .033 |
|  |  |  | -13 | -75 | 49 | 2.62 | .034 |
| Left | Secondary somatosensory cortex/Parietal operculum | 89 | -47 | -17 | 9 | 2.49 | .043 |
| Left | Thalamus | 58 | -19 | -23 | 7 | 2.62 | .034 |
| **Negative correlation** |  |  |  |  |  |  |  |
| No voxels significantly active |  |  |  |  |  |  |  |
| **1.2.2. Isolated spindles** |  |  |  |  |  |  |  |
| **Positive correlation** |  |  |  |  |  |  |  |
| No voxels significantly active |  |  |  |  |  |  |  |
| **Negative correlation** |  |  |  |  |  |  |  |
| Right | Supplementary motor area | 205 | 17 | 3 | 69 | 2.97 | .022 |
|  |  |  | 9 | 5 | 73 | 2.68 | .036 |
|  |  |  | 13 | -3 | 75 | 2.49 | .049 |
| **2. NREM3 sleep** |  |  |  |  |  |  |  |
| **2.1. Regression between functional connectivity of the hippocampus and offline performance gains** |  |  |  |  |  |  |  |
| **2.1.1. Grouped spindles** |  |  |  |  |  |  |  |
| **Positive correlation** |  |  |  |  |  |  |  |
| Right | Supramarginal gyrus | 328 | 65 | -19 | 35 | 3.04 | .019 |
|  |  |  | 65 | -19 | 41 | 2.87 | .024 |
| **Negative correlation** |  |  |  |  |  |  |  |
| No voxels significantly active |  |  |  |  |  |  |  |
| **2.1.2. Isolated spindles** |  |  |  |  |  |  |  |
| **Positive correlation** |  |  |  |  |  |  |  |
| No voxels significantly active |  |  |  |  |  |  |  |
| **Negative correlation** |  |  |  |  |  |  |  |
| Left | Primary motor cortex | 104 | -7 | -23 | 49 | 3.41 | .020 |
| Right | Precuneus | 65 | 9 | -37 | 47 | 3.05 | .034 |
| **2.2. Regression between functional connectivity of the putamen and offline performance gains** |  |  |  |  |  |  |  |
| **2.2.1. Grouped spindles** |  |  |  |  |  |  |  |
| **Positive correlation** |  |  |  |  |  |  |  |
| No voxels significantly active |  |  |  |  |  |  |  |
| **Negative correlation** |  |  |  |  |  |  |  |
| No voxels significantly active |  |  |  |  |  |  |  |
| **2.2.2. Isolated spindles** |  |  |  |  |  |  |  |
| **Positive correlation** |  |  |  |  |  |  |  |
| No voxels significantly active |  |  |  |  |  |  |  |
| **Negative correlation** |  |  |  |  |  |  |  |
| Left | Primary motor cortex | 165 | -5 | -21 | 49 | 3.51 | .016 |
|  |  |  | -3 | -19 | 53 | 3.50 | .017 |

Note: k = cluster size (determined at a threshold of $P = .05$ after small-volume correction). Minimum cluster size = 5 voxels. FWE-SVC = family-wise error small-volume correction. x, y, and z coordinates are specified in the MNI space.

